# Selective inhibition of goal-directed actions in the mesencephalic locomotor region

**DOI:** 10.1101/2022.01.18.476772

**Authors:** Nadine K Gut, Duygu Yilmaz, Krishnakanth Kondabolu, Icnelia Huerta-Ocampo, Juan Mena-Segovia

**Affiliations:** Center for Molecular and Behavioral Neuroscience, Rutgers University; Newark, NJ, USA

**Keywords:** pedunculopontine, dopamine, willing action, bradykinesia, Parkinson’s disease, basal ganglia, dlight dopamine sensor, motor, action selection, goal-directed

## Abstract

Dopamine enables purposive behavior and adjusts vigor as a function of the relative value of actions. In Parkinson’s disease, dopamine neurons die and give rise to a series of motor and cognitive changes that interfere with the expression of volitional actions. Here we report a novel inhibitory input to dopamine neurons originated in the mesencephalic locomotor region that selectively blocks purposive behavior. GABAergic neurons of the pedunculopontine nucleus (PPN) synapse onto dopamine neurons of the substantia nigra and decrease dopamine release in the dorsal striatum. Activation of PPN neurons abolished exploratory locomotion and goal-directed actions while preserved other motor behaviors; furthermore, PPN caused a decrease in movement vigor and interrupted motor sequences presumably by modulating the immediate value of the learned action. Our results reveal an inhibitory mechanism in the midbrain that rapidly and reversibly adjusts the intrinsic value of ongoing actions.

## INTRODUCTION

Successful interaction of animals with their constantly changing environment requires ongoing decision-making with the aim of selecting the behavioral response that results more profitable in a certain context. A key aspect in this selection process is the evaluation of the *utility* of a range of possible actions from the behavioral repertoire based on the subjective valuation of an outcome (Shadmehr et al., 2019). Thus, actions that lead to increased value of the outcome (i.e. large rewards) have increased utility over actions with minimal value (i.e. small or no rewards) (Kawagoe et al., 1998; Seideman et al., 2018; Summerside et al., 2018). Enhanced utility therefore results in an increased probability of any given action being selected and adjusted for effort expenditure (i.e. gain in vigor). How animals perceive the value of and assign utility to their selected actions and consequently engage in purposive behavior is not well understood.

Dopamine is essential for the generation of purposive behavior. Dopamine neurons encode the value of behavioral outcomes and consequently have a primary role in the selection of actions (Howard et al., 2017). The release of dopamine in the striatum, the main input structure of the basal ganglia, is considered to regulate movement initiation, movement kinematics, and the encoding of reward prediction error (Bayer and Glimcher, 2005; Coddington and Dudman, 2018; Cohen et al., 2012; Engelhard et al., 2019; Hamid et al., 2016; Schultz et al., 1997; da Silva et al., 2018). Critically, dopamine responses encode reward value *during* the execution of an action sequence that leads to a reward (Jin and Costa, 2010), signal the distance and size of the rewards (Howe et al., 2013) and encode reward expectation (Coddington and Dudman, 2018). Dopamine transmission has been further associated with the initiation and termination of self-paced action sequences (Jin and Costa, 2010), and the tuning of vigor during voluntary purposive movements (Panigrahi et al., 2015). Accordingly, loss of dopamine activity directly impinges on the expression and vigor of volitional actions (Panigrahi et al., 2015; da Silva et al., 2018), as seen in Parkinson’s disease (Ferrazzoli et al., 2018; Mazzoni et al., 2012). In sum, dopamine activity dynamically updates the perceived value and therefore the utility of actions during their execution as a mechanism to modulate the gain of purposive behavior.

Essential for understanding how dopamine assigns utility during motor performance is the characterization of the excitatory and inhibitory modulation of dopamine neurons. One of the major sources of innervation to dopamine neurons arises in the mesencephalic locomotor region (MLR), which comprises a mixture of cholinergic, glutamatergic and GABAergic neurons. Excitatory neurons of the MLR modulate a range of locomotor responses and movement preparation, but less is known about its GABAergic population. The pedunculopontine nucleus (PPN) has recently received significant attention due to its role as part of the MLR (Caggiano et al., 2018; Dautan et al., 2021; Josset et al., 2018) and its capability to target functionally distinct subsets of dopamine neurons, switch the firing mode of dopamine neurons, and modulate reinforcement and locomotion (Dautan et al., 2016; Xiao et al., 2016; Yau et al., 2016). PPN neurons that monosynaptically innervate dopamine neurons display functionally heterogeneous responses (Tian et al., 2016), including an excitatory response to conditioned stimuli that is compatible with the function of cholinergic neurons (Dautan et al., 2016), and an inhibitory response to reward expectation that cannot be explained on the basis of the current model of connectivity between PPN and the dopamine midbrain. Here we report the existence of a dense GABAergic projection originated in the PPN that directly synapses onto dopamine neurons of the substantia nigra. Activation of PPN GABAergic synapses on dopamine neurons modulate dopamine release in the striatum and selectively suppresses purposive behavior while leaving other motor behaviors intact. The transitory inhibition of value-guided behavior suggests that PPN GABAergic neurons shape the utility of ongoing actions through the tuning of dopamine release, impinging on the willingness of animals to persist in their intended behavior.

## RESULTS

### Dopamine neurons receive inhibitory input from PPN

To characterize the connectivity of GABAergic neurons of the PPN (hereafter PPN_GABA_), we transduced them with a double-floxed virus encoding the expression of the yellow fluorescent protein (YFP) in the cell bodies and their processes, and the red fluorophore mRuby associated to the presynaptic protein synaptophysin (**Figure 1A**). We identified axonal innervation and mRuby-positive puncta primarily in the substantia nigra pars compacta (SNc) and to a lesser degree in the ventral tegmental area (VTA); other structures included the extended amygdala, dorsal raphe, hypothalamus, thalamus, superior colliculus and lower brainstem (**Figure S1**). The density of innervation in dopamine areas, however, was at least one order of magnitude denser than in any other structure and was clearly concentrated in tyrosine hydroxylase (TH)-rich regions of the midbrain (**Figure 1B**). Quantification of the density of synaptophysin-labeled puncta in the dopamine midbrain (i.e. SNc and VTA) showed a topographical distribution where lateral portions receive more PPN_GABA_ innervation than medial portions (t_3_=6.572, two-tailed paired t-test, P=0.007, n=4; **Figure 1C, right**), thus revealing a preferential innervation of the SNc. Immunolabeling for the postsynaptic protein gephyrin, which is expressed at inhibitory synapses, revealed their appositions with synaptophysin puncta on TH-positive processes (**Figure 1D**), thus suggesting synaptic contacts. To unequivocally demonstrate the presence of synapses and characterize their morphology, we quantified the incidence of synapses in PPN_GABA_ axons onto TH processes by converting the YFP signal to a permanent peroxidase reaction and immunolabeling for TH processes with tetramethylbenzidine. PPN_GABA_ axons were observed to make synaptic contacts preferentially with identified TH-positive postsynaptic structures (59%; n=3 mice), which were identified as both, Gray’s type I (symmetric) and type II (asymmetric; 42.06% and 57.9%, respectively; **Figure 1E,F**). These results demonstrate the existence of a GABAergic projection arising in the PPN and giving rise to monosynaptic contacts with dopamine neurons.

**Figure 1.**
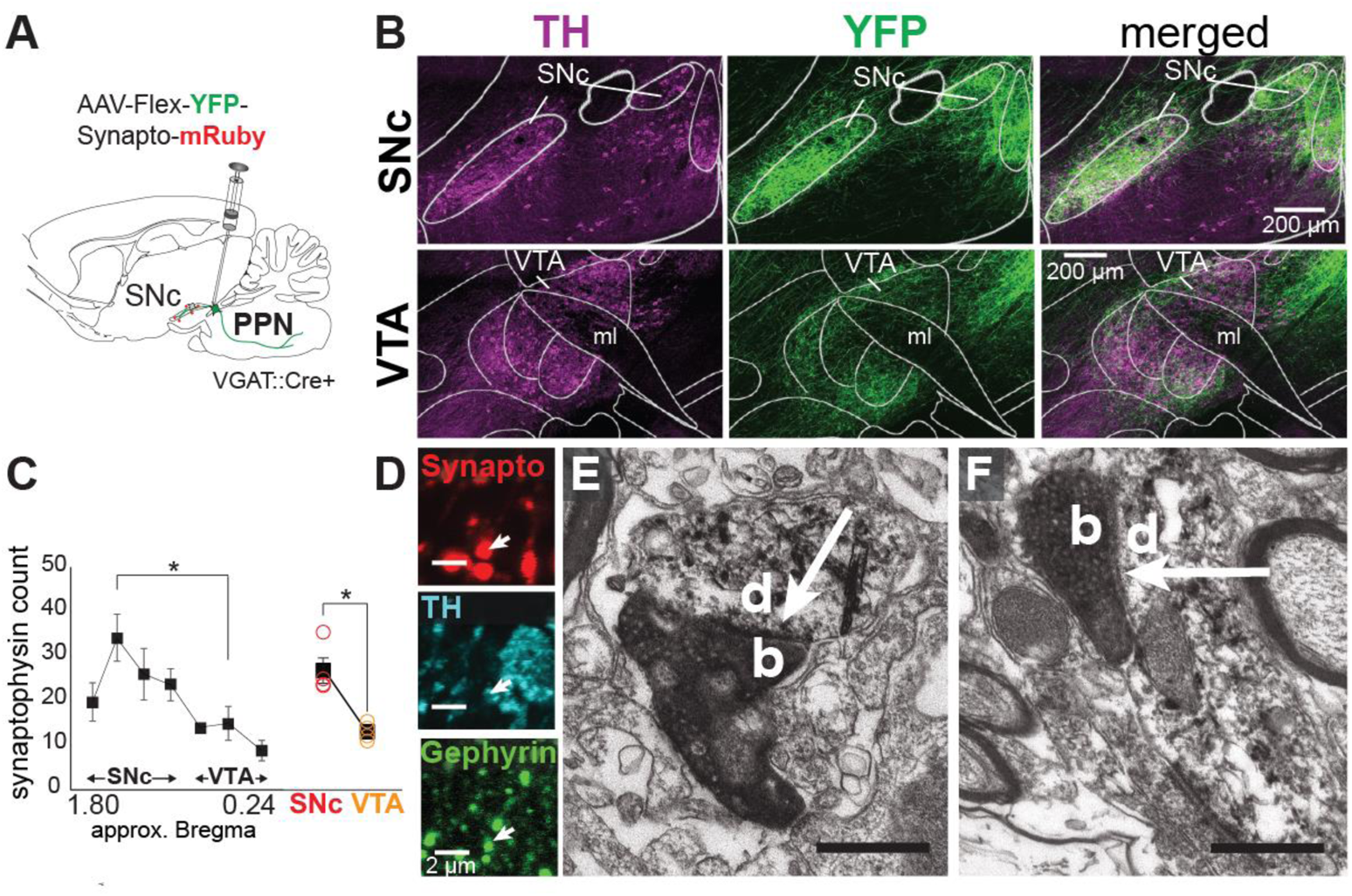
PPN_GABA_ neurons synapse onto midbrain dopamine neurons. **A**, Experimental schematic for anatomical experiments. **B**, PPN_GABA_ axons were mostly restricted to TH-rich areas. **C,** Synaptophysin quantification showed preferential distribution in lateral sections (RM-ANOVA, F_6,18_= 5.095, P=0.003; Bonferroni corrected: P=0.030), and a significantly higher density in the SNc compared to VTA. **D**, Synaptophysin-positive PPN puncta (synapto) were apposed to TH-positive processes co-expressing gephyrin. **E-F**, Electron micrographs of PPN_GABA_ boutons (b) forming asymmetric (**E**) and symmetric synapses (**F**) with TH-positive dendrites (d).

To test the effects of PPN_GABA_ axons on dopamine neurons, we transduced PPN_GABA_ neurons with channelrhodopsin-2 (ChR2) in VGAT::Cre mice and performed *ex vivo* whole-cell recordings of SNc dopamine neurons (**Figure 2A top**). Dopamine neurons were identified by the characteristic I_H_ response upon injection of current steps (**Figure 2A bottom**) and responded with inhibitory postsynaptic currents (IPSCs) of 36.31±9.37 pA and with a latency of 3.88ms±0.31ms when exposed to a 2ms-long pulse of blue light; in current-clamp settings, the average postsynaptic response was −3.14±0.67 mV (**Figure 2C-E**). These responses were abolished by bicuculline, a GABA-A receptor antagonist (**Figure 2F**; two-tailed paired t-test, t_6_=3.77; P=0.009, n=7), and restored once the blocker was washed out (**Figure 2B)**. This experiment proved that PPN_GABA_ neurons monosynaptically inhibit dopamine neurons through GABA-A receptors.

**Figure 2.**
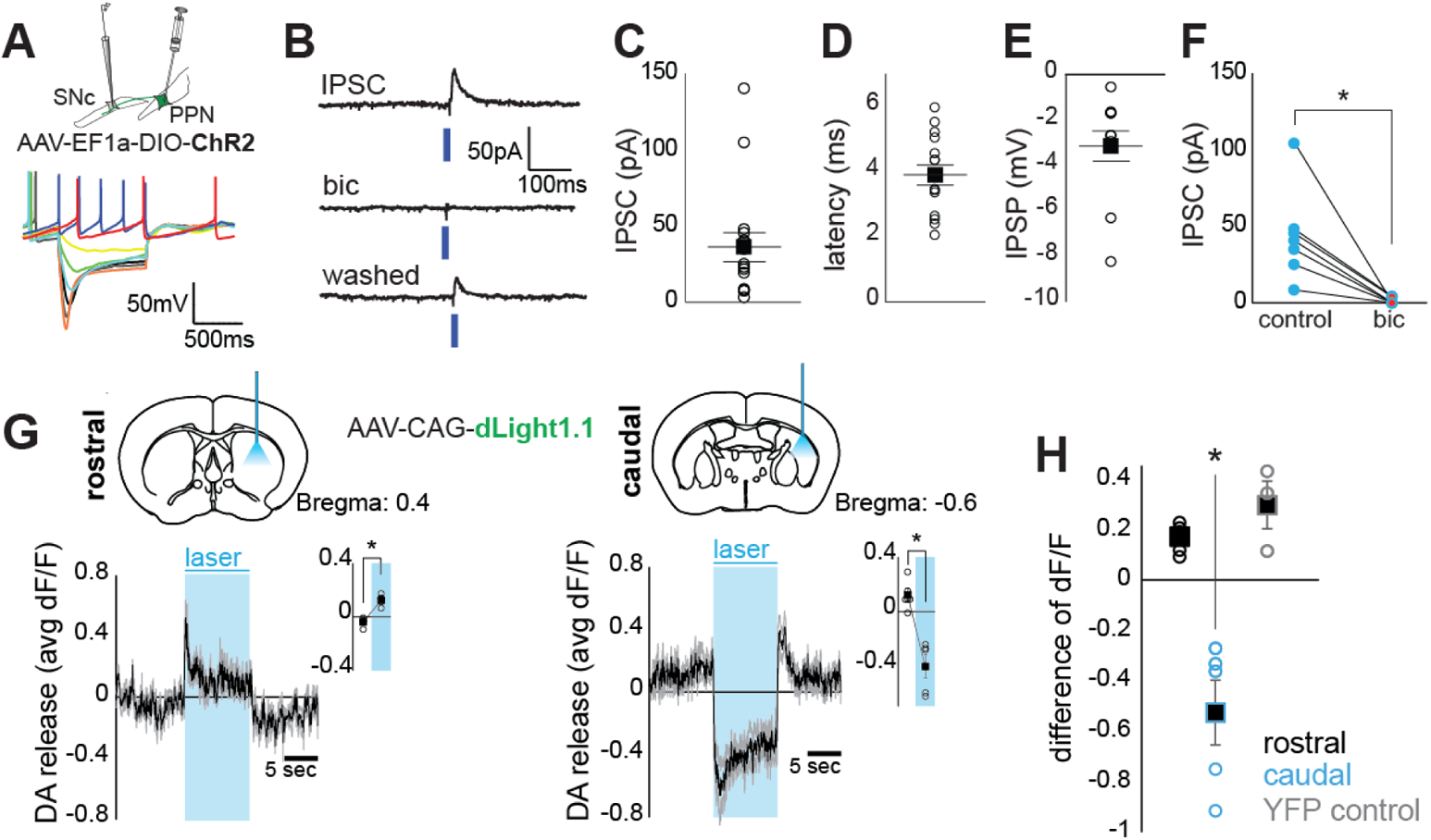
PPN_GABA_ neurons inhibit dopamine activity and decrease dopamine release in the striatum. **A**, Whole-cell recordings of dopamine neurons displaying their characteristic I_H_-current and experimental schematic. **B,** Excitation of ChR2 in PPN_GABA_ axons triggered IPSCs, which were blocked by bicuculline (bic). **C-F**, Group data showing dopamine neuron responses in voltage clamp, their latency, in current clamp and the significant reduction in current during bicuculine administration. **G,** Top, experimental schematic of striatal dopamine measurements with dLight. Bottom, average fluorescent intensity in the caudal striatum decreased significantly during PPN_GABA_ stimulation (right; insert, RM-ANOVA: F_1,4_=16.765; P=0.015; n=5), but not in the rostral striatum (left; insert, RM-ANOVA: F_1,4_=40.350; P=0.003; n=5). **H,** Compared to controls (YFP expression), PPN_GABA_ effects were significantly different in the caudal group (F_1,6_=19.834; P=0.004; n=8) but not in the rostral group (F_1,6_=2.934; P=0.138; n=8).

### PPN-mediated inhibition decreases dopamine release in the striatum

SNc neurons provide profuse dopamine innervation of the dorsal striatum. Dopamine neurons of the lateral SNc receive dense PPN_GABA_ afferents and preferentially target the dorsolateral striatum and the tail of the striatum (Menegas et al., 2018; Poulin et al., 2018). To determine the extent of inhibition of PPN_GABA_ neurons on dopamine function, we measured dopamine release in the striatum during optogenetic activation of PPN_GABA_ axon terminals in the SNc. For this purpose, we expressed ChR2 in PPN_GABA_ neurons and implanted optic fibers above the SNc. In addition, we expressed the dopamine sensor dLight (Patriarchi et al., 2018) at two distinct sites of the dorsal striatum (rostral and caudal) and implanted optic fibers for photometric recordings (**Figure 2G**). Optogenetic stimulation caused a significant decrease of the average change in fluorescence (dF/F) in the caudal striatum (**Figure 2G, right**) but not in the rostral striatum (**Figure 2G, left**; RM-ANOVA, F_1,4_=28.318; P=0.006). Changes in fluorescence in the rostral striatum did not differ from YFP controls (no ChR2; **Figure 2H, also see Figure S2**). These results show that activation of PPN_GABA_ inputs to the SNc decreases dopamine release in the caudal dorsal striatum.

### PPN_GABA_ neurons block dopamine-mediated exploratory behavior

Dopamine neurons of the lateral SNc are involved in the modulation of motor behavior through the release of dopamine in the striatum (Howe and Dombeck, 2016). To test whether dopamine neuron inhibition by PPN_GABA_ axons in the SNc has an effect on motor activity, we expressed ChR2 in PPN_GABA_ neurons and implanted optic fibers above the SNc (**Figure 3A**). We first tested the animals in the open field and observed that activation of PPN axons abolished locomotor activity (**Figure 3B-D**; two-tailed t-test t_24_=9.279, P<0.001) and blocked movement initiation (**Figure 3E**). Before stopping, mice showed shuffling of steps and decreased stride length. Following laser offset, mice had difficulties to reinitiate locomotion, showing a variable delay (typically ~4s; **Figure 3B**). Importantly, the episodes of locomotor arrest did not involve canonical freezing responses (Anagnostaras et al., 2010), as mice switched to other motor behaviors during the stimulation when locomotor activity was compromised, such as sniffing, grooming and rearing (**Figure 3F-H**). Similar responses were observed irrespective of the duration of the stimulation, where single 150-ms or 300-ms pulses significantly reduced speed for 5s (150ms-pulses data not shown; **Figure 3I-K**; two-tailed t-test, t_12_=6.596; P<0.001, n=14). These results suggest that a dopamine-mediated shift in the striatal output takes place during PPN activation possibly causing disinhibition of D2-expressing indirect spiny projection neurons (iSPNs) (Kravitz et al., 2010), as previously seen in photometric recordings of iSPNs during similar motor behaviors (Markowitz et al., 2018), leading to a marked reduction in movement speed and a total stop of previous locomotor behavior. To test whether this locomotor effect is selective of PPN_GABA_ axon terminals in the SNc, using the synaptophysin mapping reported above (**Figure S1**), we identified two other structures with above-average presence of PPN_GABA_ puncta and which function is directly or indirectly associated with motor behavior: pontine nucleus caudalis and extended amygdala. Stimulation of PPN_GABA_ axons in these structures did not affect exploratory locomotion in the open field (**Figure S3**), thus also ruling out an antidromic effect mediated by PPN_GABA_ axon collaterals.

**Figure 3.**
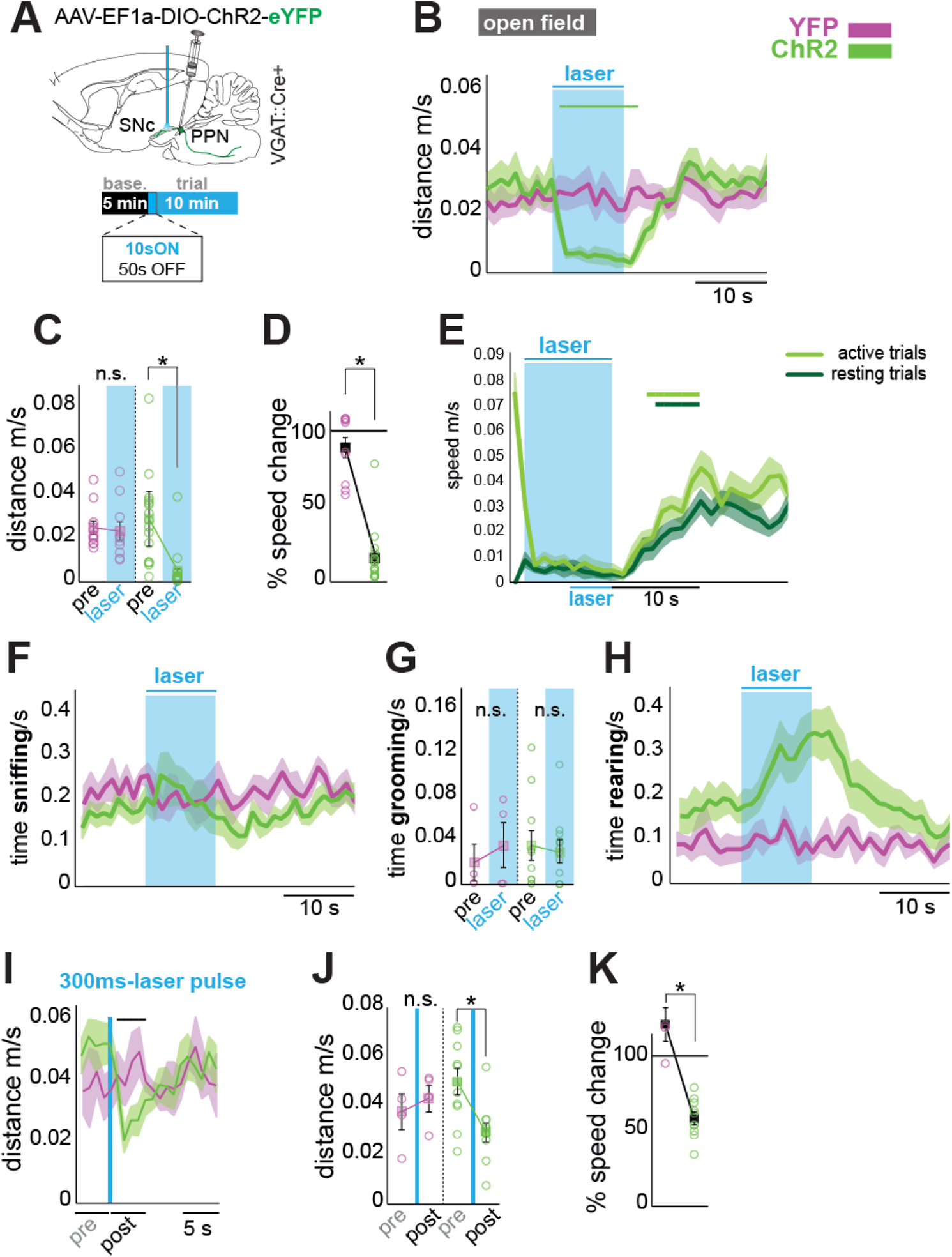
PPN_GABA_ neurons block dopamine-mediated exploratory behavior. **A**, Experimental schematic for motor experiments. Optic fibers were implanted above the SNc. **B,** Locomotor activity (m/s) was significantly reduced during PPN_GABA_ stimulation (trial average, 10s-on, 50s-off, 20ms pulses at 20Hz; mixed ANOVA: F_15,360_=5.509; P<0.001; main effect of time in ChR2 mice: F_15,225_=14.661; P<0.001, main effect of time in YFP control mice: F_15,135_=0.819; P=0.6550). **C,** Total distance traveled was reduced significantly during light delivery only in ChR2-transduced mice (mixed ANOVA: F_1,24_=15.397, n=26, P=0.001; laser main effect, ChR2: F_1,15_=22.595, P<0.001; control: F_1,9_=1.241; P=0.294). **D,** Speed was significantly reduced in ChR2 mice compared to controls. **E,** To detect potential variability in the locomotor effects following PPN_GABA_ stimulation dependent on the behavioral state of the animal, trials of open field stimulation were separated according to whether mice were moving (active trials) or not (resting trials) in the 5s that preceded the laser delivery. No difference was observed between the two types of trials (two-way RM-ANOVA, F_10,150_=1.598; P=0.169), suggesting that PPN stimulation blocks movement initiation in resting trials. Mice moved significantly faster 5-6s after laser offset (RM-ANOVA: ChR2_[resting]_: F_10,150_=14.136; P<0.001, post-hoc (Bonferroni corrected): 6th second: P=0.007; RM-ANOVA: ChR2_[active]_: F_10,150_=19.288; P<0.001, post-hoc (Bonferroni corrected): 5th second: P=0.000; n=16). **F-H,** Other motor behaviors in the open field were not abolished, such as sniffing (P=0.075), grooming (P=0.480) and rearing, which increased during the stimulation (RM-ANOVA: F_10,120_=2.258; P=0.019; main effect of time for ChR2 mice: F_10,90_=5.581; P<0.001 n=10). **I-K,** Single 300ms laser pulses significantly reduced locomotor activity, as shown during trial average (**I,** mixed ANOVA, F_10,120_=4.043; P<0.00; time main effect [RM-ANOVA], ChR2: F_10,90_=9.384, P<0.00; control: F_10,30_=0.668, P=0.744), total distance traveled (**J,** mixed ANOVA: F_1,12_=24.901; P<0.001; laser main effect, ChR2: F_1,9_= 44.918; P<0.00; control: F_1,3_=3.970, P=0.140) and change of speed (**K**).

### PPN_GABA_ neurons do not elicit behavioral arrest

To determine the extent of the motor effects produced by PPN_GABA_ inputs to dopamine neurons, we next trained the animals in the rotating rod task (RotaRod) and tested their locomotor capability during accelerating speeds. Strikingly, optogenetic activation of PPN_GABA_ axons did not interfere with locomotor activity in this task, as experimental mice remained walking with fluid strides in the rod for as long as control mice did (**Figure 4A**; Mixed ANOVA: F_1,12_=1.745; P=0.221). Next, to test motor control in a non-locomotor task, we trained mice to keep hold of a vertical pole, re-orient their heads looking down and walk it down in order to avoid falling off. Despite PPN_GABA_ axon stimulation, mice were able to complete the task with no differences between experimental and control groups during re-orientation or transit (**Figure 4B**). To further dissect the motor effects, we tested whether PPN-driven inhibition was capable of blocking innate motor behavior. We challenged the mice to the pasta handling task, where animals are required to use their upper limbs to hold a piece of pasta, bring it to their mouths and eat it, thus testing execution of fine action sequences and feeding behavior. Neither of these behaviors, nor eating velocity, showed any difference between PPN_GABA_ axon-stimulated mice and controls (**Figure 4C**). Finally, because dopamine release is involved in positive reinforcement, we asked whether the PPN_GABA_-driven decrease in striatal dopamine triggered an aversive effect that could explain the locomotor inhibition. Mice were trained in a conditioned place aversion task and ChR2 was stimulated in one of the two chambers for 2 days. Training sessions showed an increase in time spent in the stimulation chamber as a result of impaired locomotion, similar to the effect observed in the open field. However, on the fourth day, when mice were tested on their preference for either chamber, they spent equal amounts of time in each chamber, indicating that the effect of the stimulation is not aversive (**Figure 4D-F)**. These results show that inhibition of dopamine neurons by PPN_GABA_ axons does not produce an overall motor effect nor aversion. Contrasting these findings with the open field data, altogether the results reveal a selective effect on exploratory behavior.

**Figure 4.**
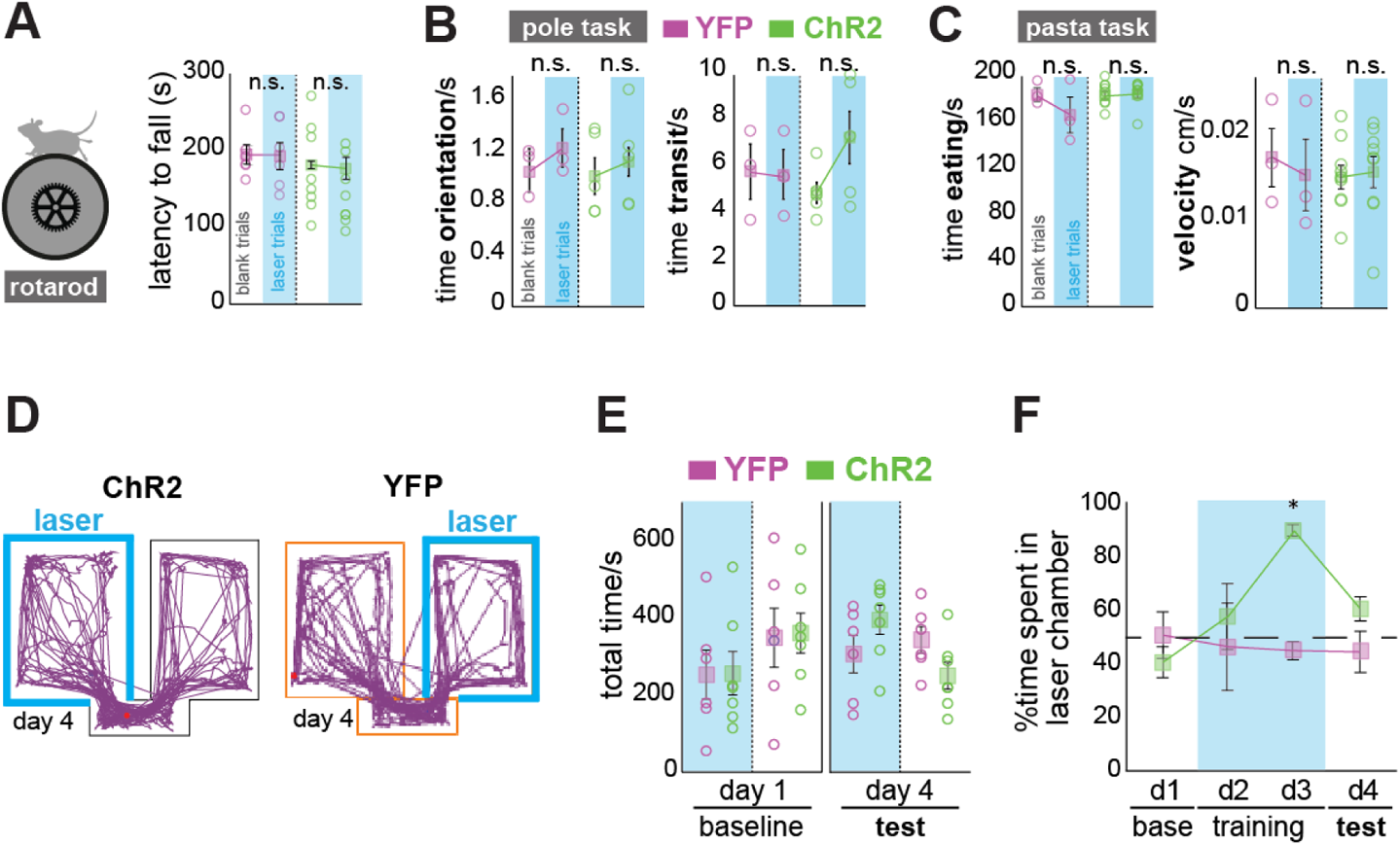
PPN_GABA_ neurons do not elicit behavioral arrest and do not produce aversive conditioning despite the drop in dopamine release. **A,** Despite the locomotor effects observed in the open field, PPN_GABA_ stimulation did not affect the locomotor performance during the Rotatod task. **B,** Control and ChR2 mice solved the vertical pole task with comparable efficiency: orientation time was not reduced during stimulation (mixed ANOVA: F_1,6_=0.076; P=0.791), neither was transit time to reach the ground (mixed ANOVA: F_1,6_=1.492; P=0.268; ChR2 n=5; control n=3). **C,** The ability to handle, manipulate and eat dry pasta was not affected either. Control and ChR2 mice ate for a comparable amount of time (mixed ANOVA: F_2,20_=3.625; P=0.045; ChR2 main effect (blank/laser): F_2,16_=1.108; P=0.354; controls main effect (blank/laser): F_2,4_=1.616; P=0.306) and had similar eating velocities (mixed ANOVA: F_1,10_=1.186; P=0.302; ChR2 n=9; control n=3). **D,** Representative activity traces of an experimental and a control mouse in the 2-chambers conditioned place aversion apparatus during the testing day (day 4). **E,** On day 1 (baseline) and day 4 (test, no laser) the total time spent in either chamber did not differ (three-way mixed ANOVA: F_1,11_=0.671, P=0.430; 2-way interaction: day*group: F_1,11_=0.03, P=0.867; laser*group: F_1,11_=3.295, P=0.097). **F,** Time spent in the laser-paired chambers was increased in ChR2 mice compared to YFP mice during laser stimulation (F_3,33_=3.207, P= 0.036; simple main effect of group on day2: F_1,11_=0.329, P=0.578; simple main effect of group on day 3: F_1,11_=139.353; P<0.001; ChR2 n=8, control n=5).

### PPN_GABA_ neurons interrupt the execution of goal-directed action sequences

Exploratory behavior is necessary to develop goal-directed actions as animals aim to maximize the reward by gaining information about their environment. In the above experiments, purposeful motor behavior could not be evaluated due to the absence of an incentive that establishes value, and thus the open field test offers restricted interpretations. To test whether mice can perform purposive goal-directed actions despite their exploratory capabilities being reduced by PPN inhibition, we trained mice to perform a stereotyped action sequence leading to reward (therefore assigning utility to the learned action sequence). Mice were trained on a self-paced operant task in which an uninterrupted sequence of eight lever presses led to a sucrose reward (see methods for details (Tecuapetla et al., 2016); **Figure S4**, **Figure 5**). We then asked whether an already initiated action sequence is affected by PPN_GABA_ axon stimulation in the SNc. Mice were stimulated for 5s triggered by the first lever press (laser on 50% of trials; **Figure 5A**) producing a significant drop in both the number of presses and the average lever press rate (**Figure 5B-D**; Mixed ANOVA: F_1,7_=19.579; P=0.003; effect of laser on ChR2 mice: F_1,5_=45.436; P=0.001;). Notably, after laser offset, mice make up for the missing number of lever presses necessary for a successful sequence with a high level of accuracy (**Figure 5E**) while maintaining a similar inter-press interval, the latter suggesting a preserved level of effort post-stimulation. The total number of trials (**Figure 5F**) and number of presses (**Figure 5G**) did not change between groups nor between laser and blank (i.e. non-stimulated) trials, suggesting that motivation is not dampened by the interruption and that the structure of the motor sequence is preserved. Similar results but at a different scale were observed when we substituted the 5s stimulation trains for a single 300ms pulse (**Figure S5**); in this situation, the mice showed no stopping but rather a larger spread of lever presses than controls immediately after laser presentation. These data suggest that PPN_GABA_ neurons momentarily block the completion of ongoing goal-directed action sequences but do not interfere with the structure of the motor program if the sequence is already initiated.

**Figure 5.**
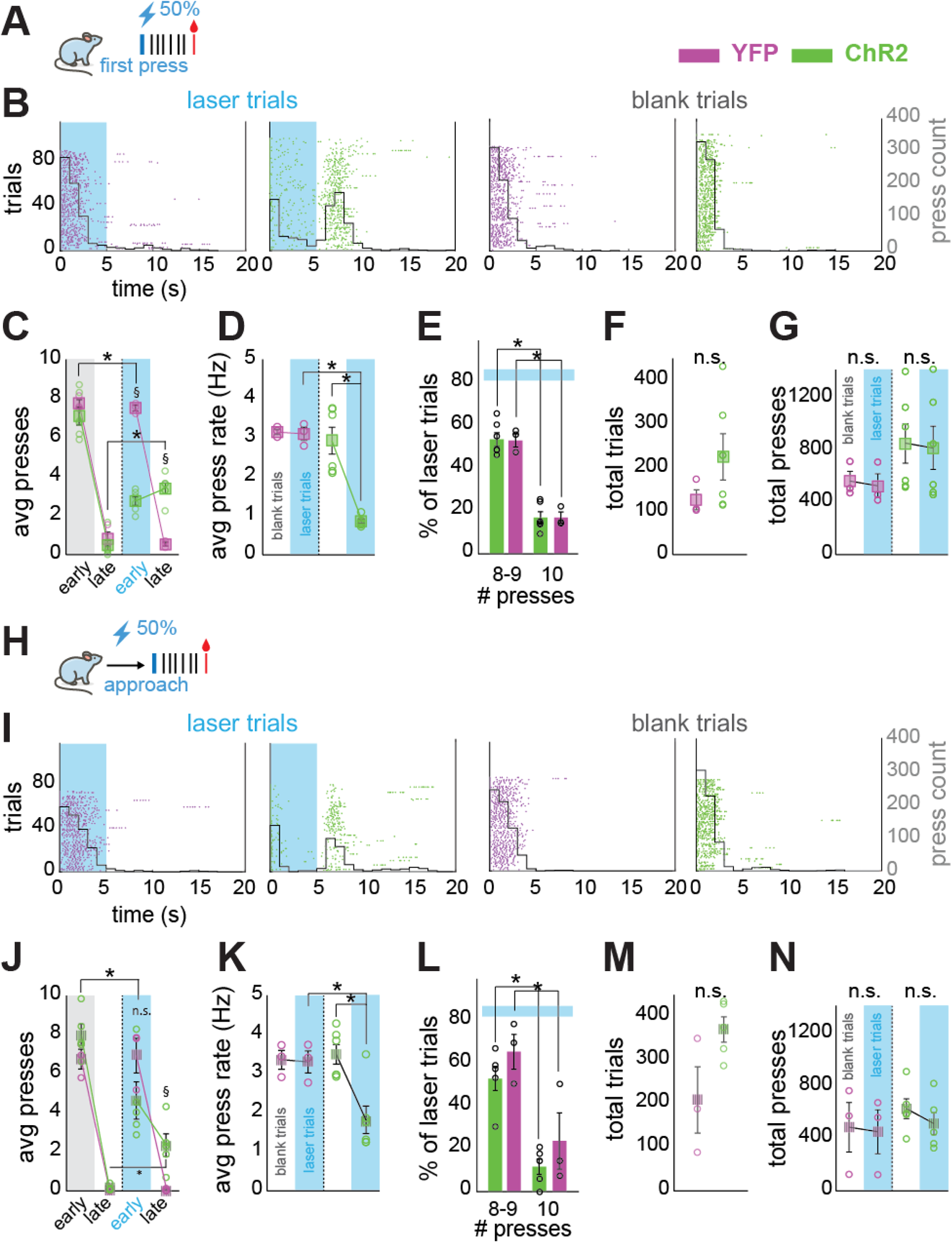
PPN_GABA_ activation impairs the execution and initiation of goal-directed action sequences. **A**, Mice were trained in an FR8 motor sequence and tested during the task execution (blue light delivered in SNc during the first lever press of the sequence in 50% of trials; 20ms pulses, 20Hz for 5s; ChR2 n=6; control n=3). **B,** Representative raster plots of stimulated (laser) and blank trials in ChR2 and controls, aligned to the first lever press. PPN_GABA_ stimulation markedly reduces the number of lever presses within each sequence during laser delivery; missing presses are completed once the stimulation is over. **C,** Lever presses were divided in early presses (*during* stimulation, 0-5s) or late presses (*after* stimulation, 5-10s) based on the distribution shown in **B**; this revealed a significant reduction in early presses and a significant increase in late presses in the ChR2 group during the laser trials (3-way mixed ANOVA: F_1,7_=42.551; P<0.00; trial-type effect [laser/blank], ChR2_[early]_: F_1,5_=96.277; P<0.00; ChR2_[late]_: F_1,5_=70.262; P<0.00 (*); group effect [control/ChR2], presses_[early]_: F_1,7_=198.834; P<0.00; presses_[late]_: F_1,7_=49.667; P<0.00 ($)). **D,** Average press rate decreased during PPN_GABA_ stimulation. **E-G**, Action sequences were completed with high accuracy, the majority of trials with 8-9 presses (Univariate ANOVA: Simple main effect of sequence length: YFP control: F_1,14_=46.658; P<0.001; ChR2: F_1,14_=95.887; P<0.001.), No effects were observed in the number of completed trials (P=0.252) nor total number of presses (P=0.971). **H,** In the following task, the same mice were then tested during the task initiation (blue light was delivered when mice crossed a proximity threshold before reaching the lever in 50% of the trials, same parameters as above; ChR2 n=5; control n=3). **I,** Raster plots as above. **J-K,** Similar to the previous experiment, ChR2 mice showed a decrease in early presses and an increase in late presses (3-way mixed ANOVA: F_1,6_=6.887, P=0.039; trial-type effect [laser/blank], ChR2_[early]_: F_1,4_=9.86; P=0.035; ChR2_[late]_: F_1,4_=14.575; P=0.019 (*); group effect [control/ChR2], presses_[late]_: F_1,6_=8.350; P=0.028), and a decrease in the press rate. **L-N,** As above, the majority of trials were completed accurately with 8-9 presses and there was an increase in the number of trials in ChR2 mice but none of these were statistically significant (**L-M**; Univariate ANOVA; Simple main effect of sequence length: YFP control: F_1,12_=12.552; P=0.004; ChR2: F_1,12_=20.614; P=0.001; and P=0.053, respectively); the total number of presses was not affected (**N,** P=0.812).

### Dopamine inhibition by PPN_GABA_ neurons blocks the initiation of action sequences

Striatal dopamine release is necessary to initiate learned motor sequences (Jankovic, 2008; Jin and Costa, 2010; Syed et al., 2016). Before sequence execution, targeted activation of iSPNs aborts motor performance (Tecuapetla et al., 2016) thus suggesting that, by means of a drop in dopamine release, PPN afferents may be capable of averting the successful initiation of goal-directed action sequences through disinhibition of D2-expressing iSPNs. To test the possibility that PPN_GABA_ neurons can block the initiation of a learned action sequence, PPN_GABA_ axon stimulation was delivered when mice crossed a proximity threshold (invisible to the animals) just before reaching the lever (**Figure 5H**). Similar to the previous experiment, mice showed a significant drop in the number of presses and the average press rate during the laser stimulation (**Figure 5I-K,** Mixed ANOVA: F_1,6_=30.701; P=0.001; effect of laser on ChR2 mice: F_1,4_=71.216; P<0.001), a similar recovery of missing presses (**Figure 5L**) while the number of trials and total presses did not change (**Figure 5M,N**). However, in contrast to the transitory interruption in the execution of the action sequence observed in the previous experiment, stimulation of PPN_GABA_ axons during action preparation significantly decreased the probability of mice completing successful sequences. Action sequences were either completely disrupted when mice checked the reward magazine before the completion of the sequence, leading to more unsuccessful trials (**Figure 6A right**; mixed ANOVA: F_1,6_=12.310; P=0.013; RM on ChR2: F_1,4_=14.288; P=0.019), or when mice aborted the action sequence (no lever press for >10s after laser onset; **Figure 6B**). Congruently, we identified an increased number of instances when mice pressed full lever press sequences (i.e. 8 or more presses) immediately after the laser offset, even in cases when they managed to make a few lever presses before stopping, suggesting that they reset the ongoing sequence and initiated new trials after laser stimulation, instead of completing an already initiated trial (data not shown). These results suggest that activation of PPN_GABA_ neurons before an action is initiated is capable of resetting the execution of a learned, goal-directed motor program.

**Figure 6.**
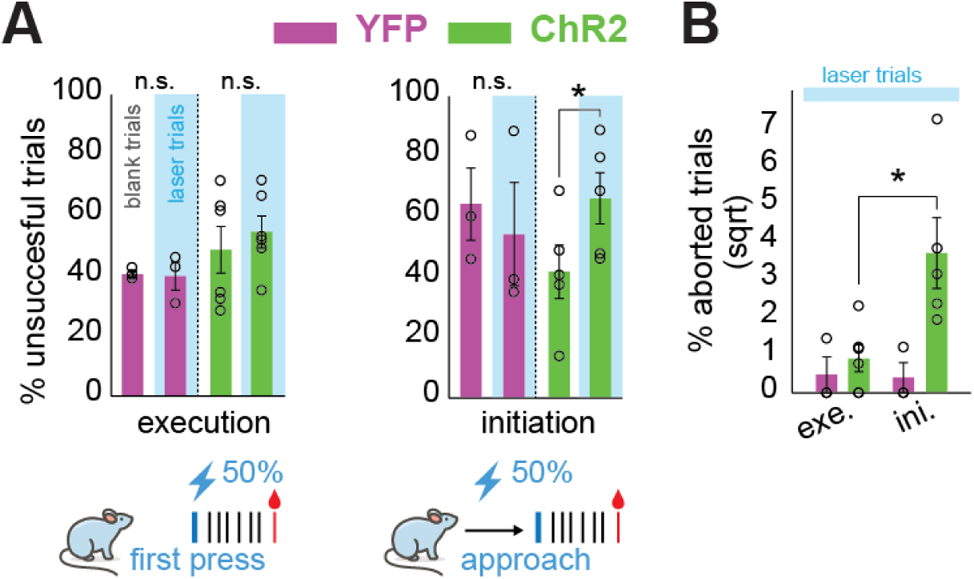
Inhibition of dopamine neurons by PPN_GABA_ axons increases the probability of aborting a motor sequence during action preparation. **A**, Optogenetic activation of PPN_GABA_ axons during action sequence initiation significantly increases the number of unsuccessful trials (i.e. due to disrupted sequence structure or abortions, see below). This effect was not observed when PPN inhibition takes place during the execution of the sequence (P=0.399). **B,** The number of aborted sequences increased significantly in the ChR2 group when the sequence initiation was disrupted by PPN inputs (mixed ANOVA: F_1,6_=7.553; P=0.033; paradigm effect [execution/initiation] in the ChR2 group: F_1,4_=17.537, P=0.014; n numbers as in Fig. 3).

## DISCUSSION

Our findings reveal the existence of a previously unknown inhibitory input to SNc dopamine neurons originated in a poorly characterized midbrain region typically associated with motor control, the mesencephalic locomotor region (MLR). We show that PPN_GABA_ neurons densely innervate the SNc and block dopamine release in the dorsal striatum, which in turn selectively blocks exploratory locomotion and purposive behavior. Once the inhibitory signal is removed, animals continue the interrupted task without displaying any reduction in the effort expended in the completion of the sequence. However, if the inhibition takes place when the animals are approaching the lever, they are significantly more likely to abort the initiated sequence. Our data further show that the inhibitory effect on dopamine neurons is capable of either obliterating purposive behavior or shifting to alternate motor states that enable default innate motor behavior (i.e. grooming, sniffing). These results suggest a role for PPN_GABA_ neurons in the gaiting of complex voluntary motor behavior.

Based on the canonical basal ganglia circuit models, the decrease in striatal dopamine levels is expected to reduce the excitability of direct pathway neurons and to disinhibit the activity of indirect pathway neurons, iSPNs. Direct pathway neurons maintain an increased neuronal discharge during entire motor sequence executions thus making them susceptible to dopamine fluctuations (Jin et al., 2014), and accordingly, dopamine release increases during lever-pressing (Syed et al., 2016). Supportive of our proposed mechanism of the decrease of vigor by the inhibition of dopamine, recent findings have shown a dopamine-dependent mechanism during which phasic activity in striatal output neurons during specific movements is sufficient to selectively reinforce changes in vigor independent of a change in motivation (Yttri and Dudman, 2016). Contrasting to the notion that structures in the lower midbrain act like output structures of the basal ganglia (Dudman and Krakauer, 2016), the results presented here thus provide the evidence of an inhibitory circuit originated in the MLR, a phylogenetically old brain structure, which is capable of radically shifting the behavioral output regulated by the basal ganglia via the control of volition.

The effects of GABA release on dopamine neurons uncovered striking similarities with motor and cognitive features in Parkinson’s disease (PD). During exploratory locomotion, while in the majority of the laser trials animals became akinetic, some trials showed instead a pronounced bradykinesia characterized by a decreased stride length and shuffling of steps, as typically seen in PD patients. Self-initiated action sequences were interrupted but not completely stopped, as some mice were still able to make occasional presses during the stimulation, however at much lower speed. These motor effects bear close resemblance with the reduction in response and movement vigor in rodent models of dopamine depletion and PD patients. In PD patients this has been portrayed as a *paralysis of the will* (Niv and Rivlin-Etzion, 2007), which describes the increased difficulty of PD patients to initiate a motor sequence when they execute internally generated actions (i.e. as opposed to external cues that trigger the initiation of the action (Mazzoni et al., 2007)). This suggests that a decrease in dopamine levels rapidly updated the intrinsic relative value of the ongoing motor sequence, reducing the vigor (as a proxy for utility) or completely abandoning the action. It then follows that adjustments in internal representations of the value of specific actions may shape the motor engagement in goal-directed actions or the intent to execute willed actions (Jahanshahi, 1998). Such adjustments depend on the encoding of value by dopamine neurons (Howard et al., 2017; Howe et al., 2013; Roesch et al., 2007) and not the motor expression of the action (Lee et al., 2019), in agreement with a role for dopamine in the facilitation of reward seeking through movement (Flagel et al., 2011; Phillips et al., 2003; Syed et al., 2016). Our results show that the expression of willed actions is contingent of the timing and intensity of opposing inhibitory signals impinging on dopamine transmission, and this is originated in the MLR. Crucially, when mice were challenged to other motor tasks (Rotarod and pole tasks), they did not show any sign of motor impairment during the stimulation. In the context of our results, the above mentioned tasks may emulate paradoxical kinesis, which occurs when PD patients are exposed to arousing or threatening situations (e.g. akinetic PD patients are able to escape a burning house (McDonald et al., 2015)). The underlying mechanisms of paradoxical kinesis are not clear yet, but it is believed that other motor systems become involved (Ballanger et al., 2006). In support of this idea, it has been shown that animals treated with dopamine antagonists were unable to move but were able to overcome motor difficulties during a stressful situation (Yntema and Korf, 1987). Our results thus reproduce some key features of dopamine function and uncover a novel regulatory mechanism that appears to have key contributions during normal and diseased conditions.

## Acknowledgements

We thank Alexandra Torres for her contribution to the analysis of anatomical data and Zirong Gu for advice on behavioral protocols. Funding: National Institutes of Health grant NS100824 (J.M.S.), NARSAD Young Investigator Award (J.M.S.) and Rutgers University.

## Author contributions

Conceptualization, JMS and NKG; Methodology: JMS, NKG, DY, KK, IHO; Investigation, NKG, DY, KK, IHO; Formal Analysis, NKG, DY, KK, IHO; Writing – original draft: NKG and JMS; Writing – reviewing and editing, JMS, NKG, DY, KK, IHO; Supervision, JMS, NKG; Funding acquisition, JMS

## METHODS

### Sample sizes, randomization and blinding

No statistical methods were used to predetermine sample size; sample sizes were rather chosen to ensure reproducibility and quantify observed effects and are similar to those generally used in the field and in our previous publications. No formal method of randomization was used, littermates were randomly assigned to different groups and there was no blinding of experimental groups. Timing of optogenetic stimulation delivery was controlled automatically.

### Mice

All procedures were performed in accordance with the Guide for the Care and Use of Laboratory Animals by the National Research Council, US, and with the approval of the Rutgers University Institutional Animal Care and Use Committee. For all experiments, all efforts were made to minimize the number of mice used.

Male and female VGAT::Cre (Jackson Laboratory, 028862) adult mice between 2 and 6 months were used for all experiments. All animals were group housed and maintained on a 12:12 light cycle (light on at 7am) with *ad libitum* access to food and water, except all behavioral animals, which were single housed and put on food restriction to slowly reduce and maintain their weight at 85% of their initial body weight. Following all surgeries, analgesics (Ketofen 2.5-5mg/kg and Buprenex SR 0.5-1mg/kg), antibiotics (Baytril, 5mg/kg or triple antibiotic ointment) and saline solution were administered.

### Viral constructs

The following Cre-dependent adeno-associated viral vectors were used in the experiments: AAV-EF1a-DIO-**EYFP**-WPRE-pA (titre: 4.6 x 10^12^, source: UNC Vector Core), AAV-EF1a-DIO-**hChR2(H134R)-EYFP**-WPRE-pA (5.6 x 10^12^, UNC), AAVdj-hsyn-flex-GFP-**synaptophysin-mRuby** (1.5 x 10^13^, Stanford Vector Core) and AAV-CAG-**dLight1.1**(cGFP) (2.0 x 10^12^, Addgene). Viruses were stored at −80°C and kept on ice during surgery.

### Virus injection and optic fiber implantation

Stereotaxic injections were performed under deep isoflurane anesthesia (0.8-1.5% in 0.8-1.2% O2) with the animals fixed in a stereotaxic frame (Kopf). For tracing of axonal innervation and for quantification of GABAergic synapses in SNc and VTA, male and female mice were transduced unilaterally with eYFP or synaptophysin-mRuby in the rostral portion of the PPN, where GABAergic neurons are densely concentrated ((Mena-Segovia et al., 2009); from Bregma, in mm: anteroposterior [AP] −4.3, mediolateral [ML] ±1.2, dorsoventral [DV] −3.4; total volume: 40nl at a rate of 5nl/min) using a Hamilton syringe. For *ex vivo* electrophysiology experiments (unilateral transduction) and behavioral/photometry experiments (bilateral transduction) mice were transduced with ChR2 in the same coordinates and with the same volume as above, whereas control mice received eYFP instead. For photometry experiments, mice were additionally transduced with dLight1.1 in the rostral dorsolateral striatum (DLS; from Bregma, in mm: AP+0.4, ML±2.2; DV-2.2 to −2.5mm; 250nl volume at 50nl/min) in one hemisphere and into the caudal striatum (from Bregma, in mm: AP −0.6, ML±2.4; DV-2.1 to −2.4mm; 200nl volume at 50nl/min) in the contralateral hemisphere (sides counterbalanced). For behavioral experiments, mice were implanted bilaterally with 200µm-diameter optic fibers (Thorlabs) at ~200µm above the SNc (from Bregma, in mm: AP-3.1mm, ML±1.5mm; DV-3.6) and secured in place with anchor screws and dental cement. In addition, custom-made 400µm-diameter optic fibers were implanted ~200µm above the dLight injection site for photometry experiments. For electron microscopy experiments, tissue of mice transduced with ChR2-YFP from the behavioral experiments was used.

### Immunohistochemistry

Following the completion of all experiments, mice were humanely euthanized with an overdose of pentobarbital solution (Euthasol; 250mg/kg) and perfused with 0.2M phosphate buffer saline (PBS) followed by 4% paraformaldehyde in PB. 50µm-thick sections were collected on a vibrating microtome (Leica) in either sagittal or coronal sections. For all experiments, sections near the injection site and/or the implantation/recording sites were further processed. For anatomical tracing experiments, all sections containing the SNc/VTA were collected and analyzed every 300µm. For the identification of transduction sites, midbrain sections were immunolabeled to reveal choline acetyltransferase (ChAT), which outlines the PPN. To facilitate identification of injection and implantation tracks sections were stained for glial fibrillary acidic protein (GFAP).

Immunohistochemical processing was initiated by blocking sections with 10% normal donkey serum in PBS-Triton (0.1%) for 1h. Primary antibodies against ChAT (host: goat, 1:250, AB144P, Millipore), green fluorescent protein (GFP, conjugated with Alexa 488; rabbit, 1:1000, A21311, Invitrogen; or unconjugated, in rat, 1:1000, 04404, Nacalai Tesque), tyrosine hydroxylase (TH, mouse, 1:1000, T2928, Millipore), gephyrin (mouse, 1:500, 147011, Synaptic Systems) or GFAP (rabbit, 1:750, AB5804, Millipore) were incubated overnight. Secondary antibodies conjugated with different fluorophores were incubated after thorough washing in PBS (Alexa405, anti-donkey 705-475-147 and anti-mouse 715-475-151; Alexa488, anti-rat, 712-545-153; Cy5, anti-goat 705-175-147, anti-rabbit 711-175-152 and anti-mouse 123-005-024; all 1:500, Jackson Immunoresearch).

For electron microscopy processing, sections of ChR2-injected mice were incubated in a cryoprotectant solution (0.05 M phosphate buffer, 25% sucrose, 10% glycerol) overnight, then freeze-thawed in order to increase penetration of the reagents. Sections were double-immunolabeled to reveal postsynaptic TH-containing structures and presynaptic YFP-containing axons. Sections were first incubated in a biotinylated antibody against GFP (raised in goat, 1:1000, AB6658, Abcam). This was followed by incubation in avidin-biotin-peroxidase complex (ABC Elite, 1:100; PK6100, Vectorlabs) for 3–4h at RT. After washing, the sections were incubated in Tris-buffer (0.5 M, pH 8; TB) containing 0.025% diaminobenzidine solution (DAB wt/vol, Sigma) and 0.5% nickel ammonium sulfate (wt/vol, Sigma) for 20 min. The peroxidase reaction was initiated by the addition of H_2_O_2_ to a final concentration of 0.01% and allowed to continue for 7–10 min. After washing thoroughly, the sections were incubated overnight at RT in anti-TH (1:1000; T2928, Millipore), followed by incubation in anti-mouse biotinylated antibody (1:300; BA-2000,Vectorlabs) for 4-6 hrs. Sections were incubated in avidin-biotin-peroxidase complex (ABC Elite, 1:100; PK6100, Vector) for 3–4 h. After the ABC incubation, the sections were washed in 0.1M PB pH 6.0. A peroxidase reaction using tetramethylbenzidine (TMB) as the chromogen was then performed at 4°C to reveal TH-positive structures. The sections were postfixed in 1% osmium tetroxide in PB for 20 min, dehydrated, infiltrated with resin (Durcupan ACM, Fluka), mounted on slides and cured at 60 °C for 48 h.

### Histological analysis

#### Quantification of YPF axonal density in whole brain sections

After immunohistochemical processing, scans were obtained at x10 magnification using an Olympus Confocal Microscope (FV1200) at three levels, 10µm apart. The density of AAV-positive axons was quantitatively assessed off-line. Images were overlapped with templates extracted from the mouse brain atlas (Franklin, K. B. J., & Paxinos, G. (1997). The mouse brain in stereotaxic coordinates.) A corrected total axonal innervation was calculated in each brain structure using the following formula: *Integrated density – (Area of selected structure x Mean fluorescence of background reading)*. Background estimation consisted of the average of the mean gray area of 2 randomly selected areas in the same section. Additional brains were qualitatively assessed by assigning a score of 1 to 6 corresponding to the axonal density.

#### Synaptophysin quantification in SNc/VTA

To quantify synaptophysin in TH-rich structures of the midbrain and obtain the density of synaptic incidence, sagittal images were taken at a x20 magnification and prepared as described above. The average distance between sections was 0.24mm, yielding 3 sections per structure (VTA and SNc). After templating, the images were converted to black and white and a 30×30µm grid was overlaid for segmentation. A manual threshold was applied (Image J) and kept consistent across all sections for the automatic counting of fluorescent puncta. All 30×30µm grid squares within the SNc and VTA, as defined by the mouse brain atlas, were counted and averaged across single sections to determine the average synaptic density for each mediolateral level (6 levels per brain in total).

#### Electron microscopy

All sections were first examined under the light microscope to determine the extent and localization of labeling within the limits of the SNc. The selected regions were cut from the slides and re-embedded for ultrathin sectioning. Serial sections (~50 nm, at least six per grid) were cut on an EM UC6 ultramicrotome (Leica Microsystems), collected on formvar-coated single-slot grids and lead-stained for 7 min. Sections were examined under a Philips Tecnai 12 electron microscope. For the synaptic incidence analysis, YFP-positive axon terminals (25 per animal), identified by the presence of synaptic vesicles and often mitochondria, were identified by the electron dense peroxidase reaction product that adhered to the internal surface of the plasmalemma and the outer membrane of organelles. Each profile was examined in each of the six serial sections to determine whether they formed a synaptic specialization with a positive or negative TH structure. TH-positive structures were identified by the presence of TMB crystals in the cytoplasm. To determine the type of synapses formed by YFP-positive axons, a total of 20 synapses were considered per animal. For each synapse, the postsynaptic target was characterized. Synapses of the asymmetrical type (Gray’s type 1) were characterized as such by the presence of presynaptic vesicle accumulation, a thick postsynaptic density, a widened synaptic cleft, and cleft material, whereas symmetric synapses (Gray’s type 2) possessed a much less pronounced postsynaptic density. Structures that did not fulfil these criteria were not considered as synaptic junctions. The digital images were analyzed using Image J, and they were adjusted for contrast and brightness using Adobe Photoshop CS5.1 (Adobe Systems).

### *Ex vivo* electrophysiology

For *ex vivo* whole-cell recordings the following general protocol was used: adult mice were anesthetized with isoflurane and then intraperitoneally injected with 100 mg/kg of ketamine-xylazine. Mice were then transcardially perfused with ice-cold N-methyl d-glucamine (NMDG)-based solution containing (in mM): 103.0 NMDG, 2.5 KCl, 1.2 NaH2PO4, 30.0 NaHCO3, 20.0 HEPES, 25.0 dextrose, 101.0 HCl, 10.0 MgSO4, 2.0 Thiourea, 3.0 sodium pyruvate, 12.0 N-acetyl cysteine, 0.5 CaCl2 (saturated with 95% O2 and 5% CO2, pH 7.2–7.4 and an osmolarity of 300-310mOsm). After decapitation the brains were extracted from the skull in a petri dish (placed on ice), containing ice cold NMDG-based solution and placed in a Leica vibratome (VT 1200S) containing an oxygenated bath of NMDG-based solution maintained at −4 °C. Brain slices of 250 or 300μm thickness were obtained and immediately transferred to an oxygenated NMDG-based solution (maintained at 35 °C) for 5 min. For recovery, slices were transferred to an oxygenated artificial CSF (ACSF) solution bath maintained at 25 °C for at least an hour before being transferred to the recording rig. Recording pipettes with impedances between 3–4 MΩ contained internal solution composed of (in mM): 130 K-gluconate, 10 KCl, 2 MgCl2, 10 HEPES, 4 Na2ATP, 0.4 Na2GTP, pH 7.3. Alternatively, voltage clamp recordings at −70mV were performed using CsCl based internal solution (in mM): 125 CsCl, 2 MgCl2, 10 HEPES, 4 Na2ATP, 0.4 GTP. For post-hoc identification of the recorded neurons, biocytin was mixed with the pipette internal solution and revealed in a reaction with streptavidin using the immunohistochemical procedure described above. Current and voltage-clamp whole-cell recordings were performed using Axoclamp 700B amplifier (Molecular Devices) and ITC-1600 digitizer (Heka Electronik). Data acquisition was done using Axograph (www.axographx.com) software. Slices were visualized using Andor ixon camera and software (Oxford Instruments). At the end of the recordings the brain slices were post-fixed in 4% paraformaldehyde for post-hoc immunohistochemistry. For optogenetic stimulation recordings in both voltage-clamp and current-clamp settings, 10 individual traces were collected, with 10 second inter-trial interval between each trace, the traces were then averaged. During *ex vivo* slice recordings, dopamine neurons were identified by their electrophysiological properties and in some cases post hoc confirmed using anti-TH antibody and streptavidin reaction to identify the whole-cell patched neuron. Photo-stimulation of ChR2 expressing PPN terminals was performed with a blue LED using 2ms-width pulses. GABA-A receptor antagonist, bicuculline, was used to confirm if the post-synaptic responses were mediated by GABA-A receptors. At the end of the recordings, 405-streptavidin (1:500, S32351, Invitrogen) was used to reveal biocytin filled neurons.

### Behavior

#### Open field locomotion

Mice were tracked in the open field (40×40cm) over a period of 30min using ANY-maze tracking software (Stoelting Co.). Animals were not habituated to the apparatus to encourage exploratory behavior, but were habituated to the room, the experimenter and to the optic fiber attachment. Throughout optogenetics experiments all mice were connected to a split multiplex optic fiber connected to a rotary joint (Doric Lenses Inc) during laser and blank (non-laser) trials. Mice were tested in two alternating blocks consisting of 5 min baseline recording followed by 10min of testing (alternating periods of 10s laser ON and 50s laser OFF), and repeated once. Laser stimulation consisted of blue laser trains (473 nm, CrystaLaser) of 20ms light pulses at 20Hz at ~6mW. In a second experiment, single 300ms laser pulses every 20s were delivered while the animals explored the open field. For analysis of locomotor behavior, the distance traveled was extracted in 1s bins. For the analysis of non-locomotor behaviors in the open field, occurrences of grooming, sniffing and rearing were scored post-hoc using ANY-maze and analyzed in 1s time bins.

#### Conditioned place aversion

Mice were trained using a non-biased, 4-day paradigm, with a two-equal chamber apparatus (40×30cm) with different wall patterns connected by a short corridor. On day one, baseline activity was measured and preference for either box was identified while mice were attached to a split multiplex optic fiber (Thorlabs) but no stimulation was delivered. On day 2 and 3 (training phase), mice were equally free to explore both chambers, however, entry to one of the 2 chambers (sides counterbalanced between mice) was coupled to photostimulation (3s-train pulses/10s, 20ms width at 20Hz) until the mouse left the chamber. Day 4 (test day) followed the protocol of day 1. Total time spent in each chamber was calculated in order to identify any avoidance to the stimulation.

#### Rotarod test

Following open field testing, mice were trained on a rotating rod (Rotarod, Harvard Apparatus) over two days until they reach criterium of remaining in the rod for a minimum of 120s at a fixed speed of 30rpm. Subsequently, they were tested during rod acceleration (from 4 to 40rpm in 300s) in 10 trials, alternating 5 laser trials and 5 blank trials over 2 days. Stimulation was delivered at the initiation of each Rotarod laser trial (20ms, 20Hz for 3s). Time to fall off the accelerating rod was recorded.

#### Vertical pole test

Mice were tested on a modified version of the vertical pole test. Animals were trained to navigate a 50cm long vertical pole (diameter: 9mm, 3 sessions) by placing them head up on the upper end of the pole, for which mice learned to turn around and climb down (criterium: task completion in less than 20s). Optic fibers were attached but no stimulation was delivered. During testing (3 sessions), laser was activated as mice were placed onto the pole and lasted until they reached the ground (20ms, 20Hz). The average time for turning around (orientation) and climbing down the pole (transit) was calculated.

#### Pasta handling test

Habituation to uncooked pasta pieces and testing took place in a cylindrical plexiglass cage (diameter: 25 cm) following a modified protocol of this task (MacLaren et al., 2014). Mice were recorded using two cameras (Logitech C920) placed on either side of the cylinder over three 10-min sessions, whilst handling and eating up to 7 pasta pieces of equal length (18.2 cm in total). Baseline sessions were conducted with the optic fiber attached but without stimulation. During testing sessions, light stimulation was delivered for 10s every min. At the end of each session, the remaining pasta pieces were measured. Eating behavior was scored post-hoc offline. Eating speed was calculated using the total amount of pasta eaten (cm/s).

#### Goal-directed action sequence task

Training and testing for the action sequence task took place in operant boxes running custom code (Med-Associates). Each box was equipped with a food magazine, a retractable lever on one side of the magazine, and a house light. Liquid reward (~30µl of 10% sucrose solution) was delivered into the reward port with a syringe pump. Head entries were detected through an infrared beam built in the magazine. No cues indicating the successful completion of a press sequence or reward delivery were used. Events inside the boxes (i.e. lever presses, head entries, laser deliveries) were recorded with a resolution of 10ms.

##### Training

After a single habituation session (30 min), during which mice received occasional rewards, mice were trained to lever press on a continuous reinforcement schedule (CR; i.e. each lever press delivered one reward). Once they collected a minimum of 30 rewards per 45-min session, they progressed gradually through different fixed-ratio schedules (FR3 to FR8). Sessions started with the house light turning on and lever insertion to the chamber, the remainder of the session was self-paced. Head entries during action sequences (before reaching the necessary press count) led to the termination of the trial and the following lever press counted towards a new sequence. Criterion for progression to the next FR schedule was set to reach 0.5 efficiency (number of presses needed for the number of obtained rewards [e.g. on a FR8 schedule: 8 × number of obtained rewards], divided by the total number of presses made by the mouse in that session) or when receiving more than 60 rewards per session (1h). In the case of failure to reach criterion, mice were allowed to repeat that schedule until they succeeded. Once mice showed steady performance (at least 2 sessions with more than 100 rewards on FR8), they were moved to the testing phase. Two mice were excluded due to poor performance. Baseline performance was tested with optic fibers attached but without stimulation.

##### Testing conditions

Mice were tested in two distinct stimulation paradigms to evaluate whether the timing of the laser delivery affected the outcome of the test. The same mice were used for both protocols, but the testing occurred sequentially (i.e., mice had to finalize testing in one protocol before starting the second). In the first protocol, laser was triggered by the first lever press of a new sequence/trial (20ms, 20Hz for 5s, or single 300ms pulses; 2 alternating sessions each, randomized order), thus testing execution of the sequence. In the second protocol, a proximity area was previously defined using Anymaze; when mice were automatically detected to cross this area, the laser was triggered (same parameters as above), thus testing initiation of the sequence. In each session, laser stimulation trials constituted 50% of the total number of trials per session, with no more than two consecutive laser trials per session. During the other 50% of trials, mice were allowed to culminate the sequence and reward retrieval without stimulation (blank trials; no more than two consecutive blank trials per session).

##### Action Sequence Quantification

A trial was defined by a head entry and collection of reward (marking the end of a trial and beginning of a new trial). Trials with a minimum of 2 presses where included in the analysis, a single press was not considered a trial^19^. Sequence lengths and durations were computed for each individual sequence. Sequence length denotes the number of presses within that sequence, and duration refers to the time difference between the first and the last press. Press rates were calculated by dividing the number of presses by sequence duration.

### Fiber photometry

Fluctuations in the fluorescence emitted by the dopamine sensor dLight were measured using fiber photometry (Doric). Two different excitation wavelengths were recorded: 465nm for dLight1.1 activity and 405nm for an isosbestic reference signal used to correct for photo-bleaching and movement-related artifacts (Lerner et al., 2015). Optical patch cables were extensively photo-bleached to reduce autofluorescence and a lock-in modulation/demodulation procedure was used. Mice were allowed to roam freely in an open field and received photostimulation for 10s every 30s (20ms, 20Hz) for a total of 20 trials for each recording site (rostral and caudal striatum). Demodulated fluorescent traces were downsampled to 120Hz, filtered and fluorescence intensity was calculated for each time point using (Ft – F0)/F0 for each channel, where Ft represents the fluorescence intensity at time t and F0 is calculated as a running average intensity variation over a window of 1 min^33^ to correct for a slow drift of the baseline signal due to photo-bleaching. The reference signal was then subtracted from the dLight trace.

### Statistical Analysis

Significance level was set at P≤0.05. Parametric testing was used whenever possible to test differences between two or more means. Normality was tested using the Shapiro-Wilk tests and Levene’s test was conducted to assess homogeneity. Mild violations of normality and homogeneity were accepted. Severe violations were addressed by transformation using square root or natural logarithm. Details of the statistical tests are described in the Supplementary Information. Main effects and interactions were followed up by planned comparisons when found significant. Statistical tests were done using SPSS (IBM), Matlab (MathWorks) and R (R core team 2020, Ime4).

### Data availability

In vivo fiber photometry data, in vitro electrophysiology and behavioral data are available under reasonable request to the corresponding author.

### Code Availability

All code used to collect and analyze data is available under reasonable request to the corresponding author.

### Contact for reagent and resource sharing

Further details and specific requests for resources, data, code, materials or protocols used in this manuscript should be directed to and will be fulfilled by the corresponding author.

---

**Supplementary Figure 1.**
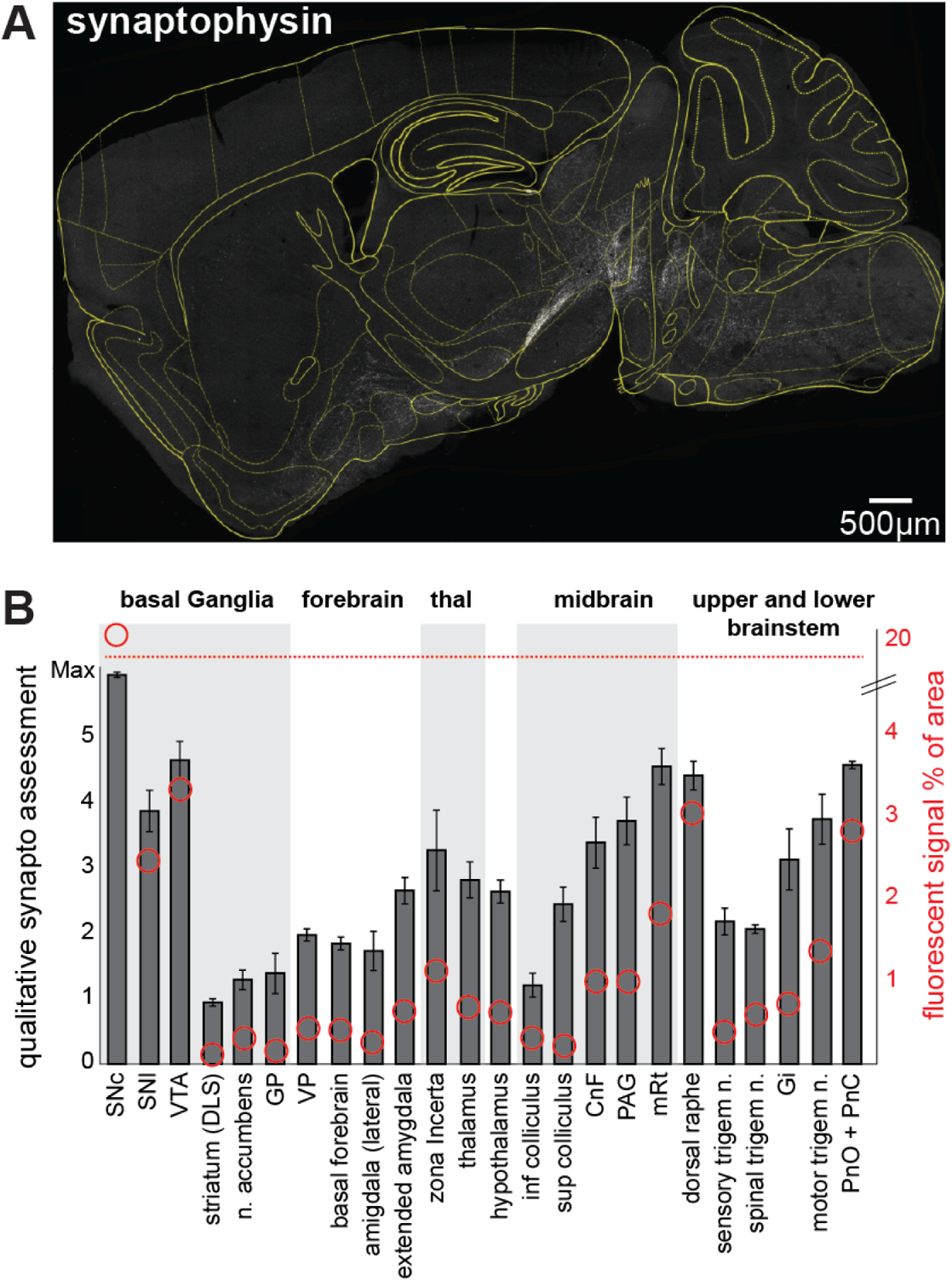
Axonal spread of PPN GABAergic neurons. **A**, Sagittal brain section at approximately 1,7mm from the midline showing fluorescent expression of PPNgaba axons after transduction with synaptophysin-mRuby (as in Fig. 1). **B,** Innervation density was first quantitatively assessed by measuring the percentage of pixels representing fluorescent signal in each selected brain structure (n=1). This initial quantification was then confirmed in 2 additional brains using a qualitative scale from 1 to 6. GP, Globus Pallidus; VP, Ventral Pallidum; CnF, Cuneiform nucleus; PAG, Periarqueductal Gray; mRt, mesencephalic reticular formation; Gi, gigantocelular reticular nucleus; PnO, Pontine reticular nucleus oral part; PnC, Pontine reticular nucleus caudal

**Supplementary Figure 2.**
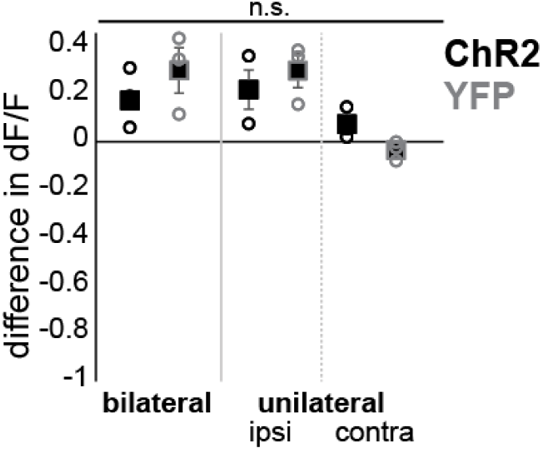
Control recordings for dopamine release measurements. Complementary dLight recordings were obtained in the rostral striatum to test possible contralateral effects of the stimulation or synergistic effects during bilateral PPN_GABA_ stimulation. No significant changes were observed across different control groups for dLight detection, including a lack of contralateral effects of PPN_GABA_ stimulation and absence of changes in control animals transduced with YFP in the PPN (mixed ANOVA: F_1,4_=1.746; P=0.235). A slight increase in fluorescence was observed during unilateral and bilateral stimulation, suggesting a possible effect of heat transduction from the stimulating laser, as reported before (Owen et al., 2019).

**Supplementary Figure 3.**
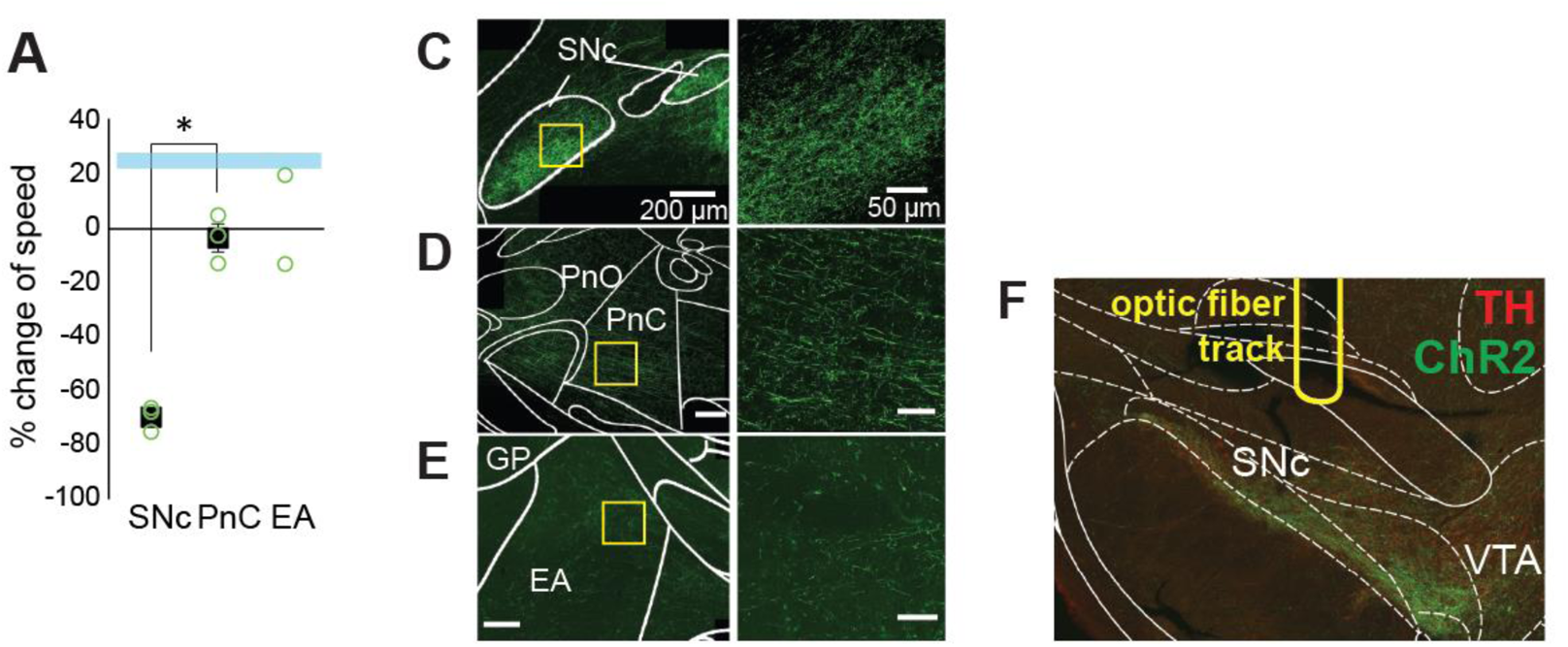
Control experiments for exploratory locomotion. **A**, Stimulation of axonal terminals in the PNc and EA did not cause a reduction of locomotion as shown when stimulating in the SNc (univariate ANOVA: F_1,4_=128.709, P<0.001). **C-E.** Images showing axonal innervation in SNc, PnC and EA. SNc: N=3, PnC: N=3, EA: N=2. **F,** Representative imaging showing track of optic fiber implanted above the SNc.

**Supplementary Figure 4.**
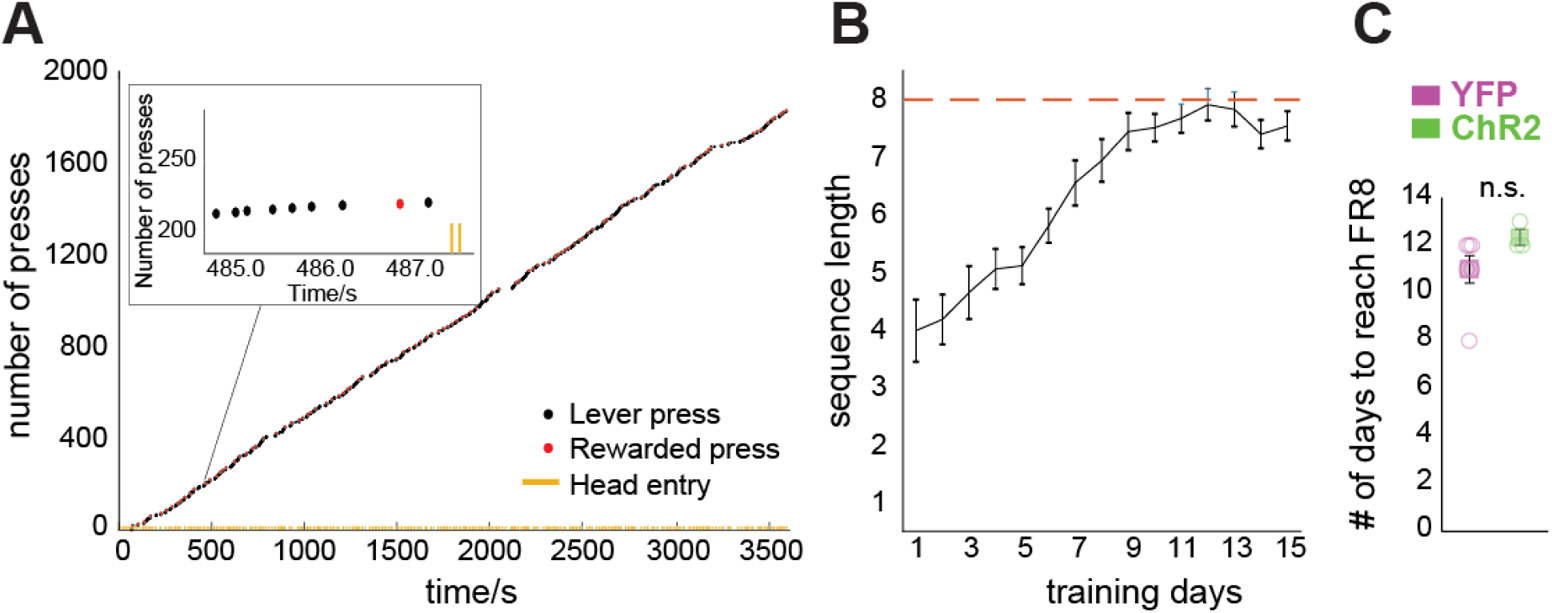
FR8 task acquisition. **A**, Representative scatterplot of the last training day before testing. Mice learned to lever press in tight bouts of 8 presses, followed by a head entry to the food magazine following the rewarded press (eighth press). **B,** Learning progression of mice. **C,** YFP (controls) and ChR2 mice needed on average 11-12 days to reach criterium for testing (minimum of 100 rewards for two consecutive days on FR8; F_1,8_=2.358; P=0.163).

**Supplementary Figure 5.**
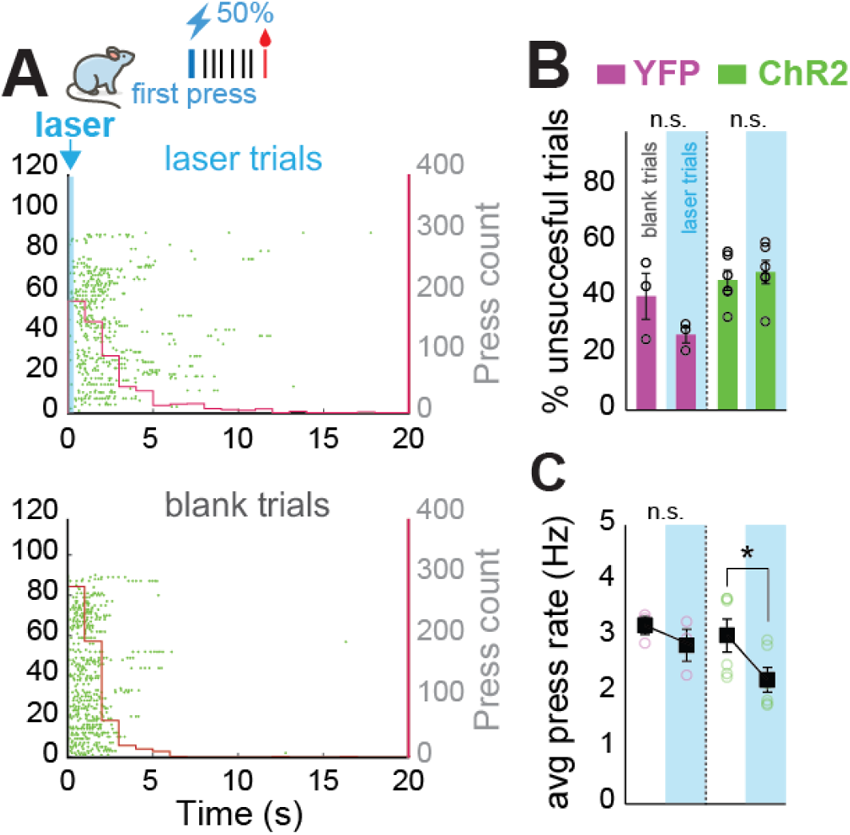
Effect of single 300ms-laser pulses on lever press performance during action sequences (FR8). **A**, Representative raster plots of an experimental mouse on laser (above) and blank (below) trials aligned to the first lever press. After a single 300ms pulse, the interval between presses increased and the presses had longer intervals. **B,** The 300ms-laser pulse had no impact on the success rate of the task (F_1,7_=282.067, P=0.005; RM-ANOVA ChR2 mice: F_1,5_=3.619, P=0.116; RM-ANOVA YFP mice: F_1,20_=6.955, P=0.119). **C,** The average press rate, however, dropped significantly (Mixed ANOVA: F_1,7_=3.207; P=0.116, effect of trial type on ChR2 mice: F_1,5_=25.752, P=0.004).

**Supplementary Table 1.** Statistical details for all figures.

| Figure | Sample size (n) | DV | Statistical approach | Exact p and critical values |
| --- | --- | --- | --- | --- |
| 1C (LHS) | DIO-Synaptophysin (4) | synaptophysin count | Repeated Measures ANOVA | $F_{6,18}=5.095$; $P=0.003$; lateral 1.5mm (SNc) vs lateral 0.54mm (VTA): **pairwise comparisons, Bonferroni corrected: $P = 0.030$** |
| 1C (RHS) | DIO-Synaptophysin (4) | synaptophysin count | two-tailed paired t-test | $t_3 = 6.572$, $P = 0.007$ |
| 2F | Chr2 (7) | IPSC (pA) | two-tailed paired t-test | $t_6=3.775$; $P=0.009$ |
| 2G | Chr2 (5) | average dF/F | 2-way Repeated Measures ANOVA | $F_{1,4}=28.318$; $P=0.006$; **Repeated Measures ANOVA for caudal (RHS): $F_{1,4}=16.765$; $P=0.015$; Repeated Measures ANOVA for rostral (LHS): $F_{1,4}=40.350$; $P=0.003$** |
| 2H | Chr2 caudal dlight (5), Chr2 rostral dlight (5) | difference of dF/F before and during laser | Repeated Measures ANOVA | $F_{1,4}=28.318$; $P=0.006$ |
| 2H | YFP control (3), Chr2 rostral dlight (5) | difference of dF/F before and during laser | Univariate ANOVA | $F_{1,6}=2.934$; $P=0.138$ |
| 2H | YFP control (3), Chr2 caudal dlight (5) | difference of dF/F before and during laser | Univariate ANOVA | $F_{1,6}=19.834$; $P=0.004$ |
| 3B | YFP control (10), Chr2 (16) | distance travelled per second | Mixed ANOVA (Greenhouse-Geisser corrected) | $F_{15,360}=5.509$; $P<0.001$; main effect of time in **Chr2 mice: $F_{15,225}=14.661$; $P<0.001$**, main effect of time in **YFP control mice: $F_{15,135}=0.819$; $P=0.655$** |
| 3C | YFP control (10), Chr2 (16) | distance travelled per second | Mixed ANOVA (Greenhouse-Geisser corrected) | $F_{1,24}=15.397$; $P=0.001$; main effect of laser in **Chr2 mice: $F_{1,15}=22.595$, $P<0.001$**; main effect of laser in **YFP control mice $F_{1,9}=1.241$, $P=0.294$** |
| 3D | YFP control (10), Chr2 (16) | %change of speed | two-tailed t-test | $t_{24}=9.279$, $P<0.001$ |
| 3E | ChR2 (16) | distance travelled per second | 2-way Repeated Measures ANOVA | $F_{10,150}=1.598$; $P=0.169$ (Greenhouse-Geisser corrected). Does timing to resume activity differ between conditions? Repeated measures for resting trials: $F_{10,150}=14.136$; $P<0.001$, **Post-hoc (Bonferroni corrected): 6th second: $P=0.007$**; Repeated measures for active trials: $F_{10,150}=19.288$; $P=0.000$, **Post-hoc (Bonferroni corrected): 5th second: $P<0.001$**. |
| 3F | YFP control (4), ChR2 (10) | time sniffing in seconds | Mixed ANOVA | $F_{10,120}=1.764$; $P=0.075$, **main effect of time for ChR2 mice: $F_{10,90}=3.008$; $P=0.003$**; main effect of time for control mice: $F_{10,30}=0.397$; $P=0.937$ |
| 3G | YFP control (4), ChR2 (10) | time grooming in seconds | Mixed ANOVA | $F_{1,12}=0.107$; $P=0.480$ |
| 3H | YFP control (4), ChR2 (10) | time rearing in seconds | Mixed ANOVA | $F_{10,120}=2.258$; $P=0.019$; **main effect of time for ChR2 mice: $F_{10,90}=5.581$; $P<0.001$** |
| 3I | YFP control (4), ChR2 (10) | distance travelled per second | Mixed ANOVA | $F_{10,120}=4.043$; $P<0.001$; main effect of time, **ChR2: $F_{10,90}=9.384$; $P<0.00$**; **YFP control: $F_{10,30}=0.668$, $P=0.744$** |
| 3J | YFP control (4), ChR2 (10) | distance travelled per second | Mixed ANOVA | $F_{1,12}=24.901$; $P<0.001$; main effect of laser in **ChR2 mice: $F_{1,9}=44.918$; $P=0.000$**; main effect of laser in **YFP control mice: $F_{1,3}=3.970$; $P=0.140$** |
| 3K | YFP control (4), ChR2 (10) | %change of speed | two-tailed t-test | $t_{12}=6.596$; $P<0.001$ |
| 4A | YFP control (6), ChR2 (8) | latency to fall in seconds | Mixed ANOVA | $F_{1,12}=1.745$; $P=0.211$ |
| 4B LHS | YFP control (3), ChR2 (5) | time for orientations in seconds | Mixed ANOVA | $F_{1,6}=0.076$; $P=0.791$ |
| 4B RHS | YFP control (3), ChR2 (5) | time for transit in seconds | Mixed ANOVA | $F_{1,6}=1.492$; $P=0.268$ |
| 4C LHS | YFP control (3), ChR2 (9) | time eating in seconds | Mixed ANOVA | **$F_{1,10}=5.032$; $p=0.049$**; Repeated Measures for YFP control: $F_{1,2}=2.496$; $P=0.255$; Repeated Measures for ChR2: $F_{1,8}=0.044$; $P=0.840$ |
| 4C<br>RHS | YFP control (3),<br>ChR2 (9) | average eating<br>speed in<br>cm/second | Mixed ANOVA | $F_{1,10}=0.396$; $p=0.302$ |
| 4E | YFP control (5),<br>ChR2 (8) | total time spent<br>in chamber in<br>seconds | 3-way Mixed ANOVA | 3-way interaction: $F_{1,11}=0.671$;<br>$P=0.430$, 2-way interaction:<br>day*group: $F_{1,11}=0.03$; $p=0.867$;<br>laser*group: $F_{1,11}=3.295$, $p=0.097$ |
| 4F | YFP control (5),<br>ChR2 (8) | % of time<br>spend in laser<br>paired<br>chamber | Mixed ANOVA | $F_{3,33}=3.207$; $P=0.036$; Simple main<br>effect of group on day2: $F_{1,11}=0.329$,<br>$P=0.578$; Simple main effect of group<br>on day 3: $F_{1,11}=139.353$; $P=0.000$. |
| 5C | YFP control (3),<br>ChR2 (6) | average<br>number of<br>presses | 3-way Mixed ANOVA | **$F_{1,7}=42.551$; $P<0.001$**; Two-way<br>interaction of trial_type and time_bin:<br>**ChR2: $F_{1,5}=99.634$; $P<0.001$**, Simple<br>main effects: no-laser vs laser: YFP<br>control early: $F_{1,2}=0.745$, $p=0.478$,<br>**ChR2 early: $F_{1,5}=96.277$; $P<0.001$**.<br>YFP control late: $F_{1,2}=0.811$; $P=0.463$,<br>**ChR2 late: $F_{1,5}=70.262$; $P<0.001$**;<br>**Univariate ANOVA YFP control vs**<br>**ChR2 during laser: early:**<br>**$F_{1,7}=198.834$; $P<0.001$**; late:<br>**$F_{1,7}=49.667$; $P<0.001$**. |
| 5D | YFP control (3),<br>ChR2 (6) | press rate in<br>Hz | Mixed ANOVA | $F_{1,7}=19.579$; $P=0.003$, **Repeated**<br>**Measures on ChR2 $F_{1,5}=45.436$;**<br>**$P=0.001$**, Repeated Measures on YFP<br>control: $F_{1,2}=0.158$; $P=0.729$;<br>Univariate ANOVA ChR2 vs YFP<br>control: non-laser: $F_{1,7}=0.199$;<br>$P=0.669$, **laser: $F_{1,7}=280.231$;**<br>**$P<0.001$** |
| 5E | YFP control (3),<br>ChR2 (6) | % of trials<br>finished with 8-9 or 10<br>presses | 2-way ANOVA | $F_{1,14}=0.940$; $P=0.940$; Simple main<br>effect of group: 8-9 presses:<br>$F_{1,14}=0.010$; $P=0.923$; 10 presses:<br>$F_{1,14}=0.000$; $P=0.992$ **Simple main**<br>**effect of sequence length: YFP**<br>**control: $F_{1,14}=46.658$; $P<0.001$;**<br>**ChR2: $F_{1,14}=95.887$; $P<0.001$.** |
| 5F | YFP control (3),<br>ChR2 (6) | total number of<br>trials | Univariate ANOVA | $F_{1,7}=1.556$; $P=0.252$ |
| 5G | YFP control (3),<br>ChR2 (6) | total number of<br>presses | Mixed ANOVA | $F_{1,7}=0.001$; $P=0.971$ |
| 5J | YFP control (3),<br>ChR2 (5) | average<br>number of<br>presses | 3-way Mixed ANOVA | **F<sub>1,6</sub>=6.887; P=0.039:** Two-way interaction of trial_type and time_bin: **ChR2: F<sub>1,4</sub>=11.652; P=0.027**, Simple main effects: no-laser vs laser: YFP control early: F<sub>1,2</sub>=0.187, P=0.707, **ChR2 early: F<sub>1,4</sub>=9.86; P=0.035**. YFP control late: F<sub>1,2</sub>=0.134; P=0.749, **ChR2 late: F<sub>1,4</sub>=14.575; P=0.019**; **Univariate ANOVA YFP control vs ChR2 during laser: early: F<sub>1,6</sub>=2.658; P=0.154; late: F<sub>1,6</sub>=8.350; P=0.028.** |
| 5K | YFP control (3),<br>ChR2 (5) | press rate in<br>Hz | Mixed ANOVA | F<sub>1,6</sub>=30.701; P=0.001, **Repeated Measures on ChR2 F<sub>1,4</sub>=71.216; P=0.000**, Repeated Measures on YFP control: F<sub>1,2</sub>=1.426; P=0.355, Univariate ANOVA ChR2 vs YFP control: non-laser: F<sub>1,6</sub>=0.118; P=0.741, laser: **F<sub>1,6</sub>=7.313; P=0.030** |
| 5L | YFP control (3),<br>ChR2 (5) | % of trials<br>finished with 8-9 or 10<br>presses | 2-way ANOVA | F<sub>1,14</sub>=0.00; P=0.984; Simple main effect of group: 8-9 presses: F<sub>1,12</sub>=1.399; P=0.260; 10 presses: F<sub>1,12</sub>=1.331; P=0.271 **Simple main effect of sequence length: YFP control: F<sub>1,12</sub>=12.552; P=0.004; ChR2: F<sub>1,12</sub>=20.614; P=0.001.** |
| 5M | YFP control (3),<br>ChR2 (5) | total number of<br>trials | Univariate ANOVA | F<sub>1,6</sub>=5.770; P=0.053 |
| 5N | YFP control (3),<br>ChR2 (5) | total number of<br>presses | Mixed ANOVA | F<sub>1,6</sub>=2.627; P=0.156 |
| 6A<br>LHS | YFP control (3),<br>ChR2 (6) | % of<br>unsuccessful<br>trials | Mixed ANOVA | F<sub>1,7</sub>=0.807; P=0.399 |
| 6A<br>RHS | YFP control (3),<br>ChR2 (5) | % of<br>unsuccessful<br>trials | Mixed ANOVA | F<sub>1,6</sub>=12.310; P=0.013; **Repeated Measures on ChR2: F<sub>1,4</sub>=14.288; P=0.019**; Repeated Measures on YFP control: F<sub>1,2</sub>=2.405, p=0.261. |
| 6B | YFP control (3),<br>ChR2 (6) | % of trials with<br>action<br>sequence<br>starting >10s<br>after laser<br>onset | Mixed ANOVA (sqrt-<br>transformed) | $F_{1,6}=7.553$; $P=0.033$; **Simple main effect of condition (during vs after) on ChR2 mice: $F_{1,4}=17.537$; $P=0.014$** |
| Suppl<br>2 | rostral dlight<br>group: ChR2 (3),<br>YFP control (3) | difference of<br>dF/F before<br>and during<br>laser | Mixed ANOVA | $F_{1,4}=1.746$; $P=0.235$ |
| Suppl<br>3A | ChR2 SNc (3),<br>ChR2 PNc (3) | % change in<br>distance<br>travelled during<br>laser | Univariate ANOVA | $F_{1,4}=128.709$; $P<0.001$. |
| Suppl<br>4C | YFP control (3),<br>ChR2 (7) | number of<br>days | Univariate ANOVA | $F_{1,8}=2.358$; $P=0.163$ |
| Suppl<br>5B | YFP control (3),<br>ChR2 (6) | % of trials | Mixed ANOVA | **$F_{1,7}=282.067$; $P=0.005$**; Repeated Measures ANOVA in ChR2 mice: $F_{1,5}=3.619$; $P=0.116$; Repeated Measures ANOVA in YFP control mice: $F_{1,20}=6.955$; $P=0.119$ |
| Suppl<br>5C | YFP control (3),<br>ChR2 (6) | press rate in<br>Hz | Mixed ANOVA | $F_{1,7}=3.207$; $P=0.116$, **Repeated Measures ANOVA on ChR2 mice: $F_{1,5}=25.752$; $P=0.004$** |

